# Dissecting resistance mechanisms in melanoma combination therapy

**DOI:** 10.1101/041855

**Authors:** Eunjung Kim, Alexander Anderson

## Abstract

We present a compartment model that explains melanoma cell response and resistance to mono and combination therapies. Model parameters were estimated by utilizing an optimization algorithm to identify parameters that minimized the difference between predicted cell populations and experimentally measured cell numbers. The model was then validated with *in vitro* experimental data. Our simulations show that although a specific timing of the combination therapy is effective in controlling tumor cell populations over an extended period of time, the treatment eventually fails. We subsequently predict a more optimal combination therapy that incorporates an additional drug at the right moment.

## I. MELANOMA RESPONSE AND RESISTANCE TO THERAPY

Metastatic melanoma is known to be resistant to chemotherapy (chemo). During the last few years, targeted therapeutic approaches have emerged as the dominant treatment choice, mainly because they target tumor cells that harbor specific genetic mutations. However, even these targeted drugs have limited long term success in treating melanoma metastatic patients, since resistance eventually emerges. Unexpectedly, when chemotherapy and a targeted treatment (AKT inhibitor, AKTi) are given in combination to metastatic melanoma patients, long-term responses were recently observed [1]. Although little is known regarding why such combinations are more successful, we suggested one possible mechanism, specifically differential induction of autophagy by mono or combination therapy [1]. To better understand how autophagy might facilitate treatment response, we developed a mathematical model comprising of a system of ordinary differential equations that explains the dynamics of melanoma cells under mono (either chemo or AKTi alone) and combination therapy of the two [2]. The model is composed of three compartments, non-autophagy and two autophagy compartments (physiological and quiescent autophagy) based on experimental observations showing that some autophagy cells continue to maintain normal cell homeostasis [3-6]. Fig. 1 shows how these three compartments interact with one another, switching rates are altered depending on the treatment being administered.

**Figure 1.**
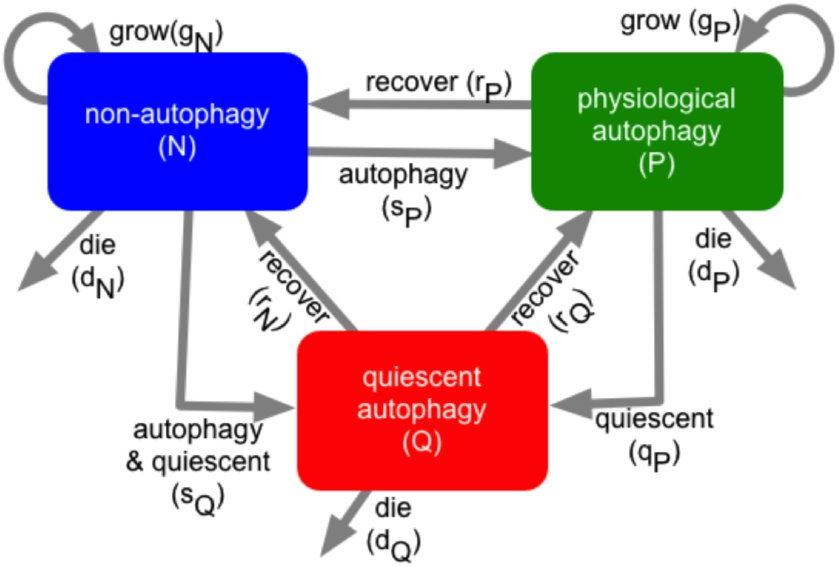
Compartmental model schematic showing how non-autophagy, quiescent and physiological autophagy interact in response to therapies.

## II. MODEL CALIBRATION AND VALIDATION

Model parameters were estimated by using the implicit filtering method to generate fitted curves describing the growth of a melanoma cell line under no treatment, mono and combination therapy (Fig. 2A). We then used the fitted model to predict the effects of 12 different schedules including no treatment, single agent therapy, concurrent combination therapy, and single agent sequential therapy of two drugs. We accurately predicted total population at day 16 in equivalent *in vitro* experiments (Fig. 2B). Interestingly, our longer-term simulation over 40 days showed that the combination therapy was effective in controlling tumor population over an extended period of time. The resistance, however, emerges (Fig. 3A) driven by the autophagy population, even for the proposed best strategy (#7 in Fig. 2B, concurrent treatment of chemo and AKTi followed by AKTi maintenance). To overcome this resistance, we applied a drug that targets the autophagy population (chloroquine (CQ)) and were able to show that additional administrations of this drug inhibited resistance (Fig. 3B).

**Figure 2.**
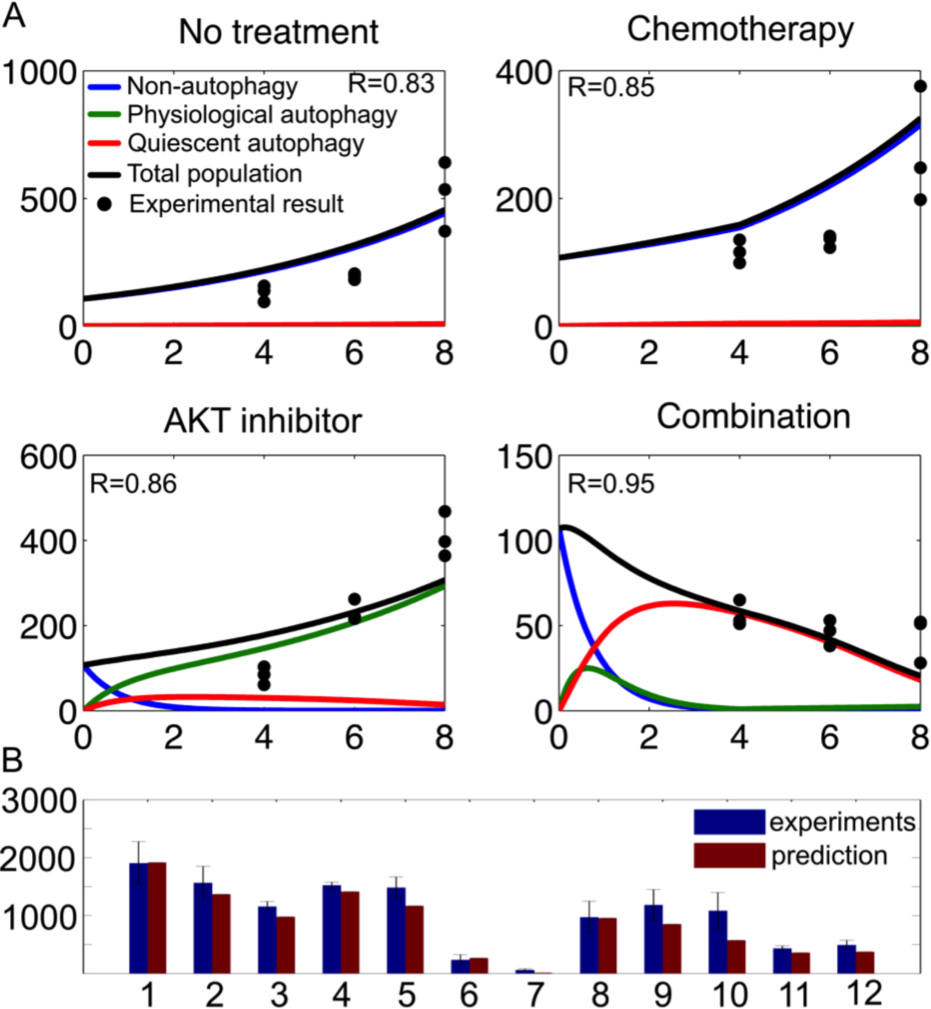
A. Calibrating the model with *in vitro* data. B. Validating model predictions with *in vitro* combination therapy.

**Figure 3.**
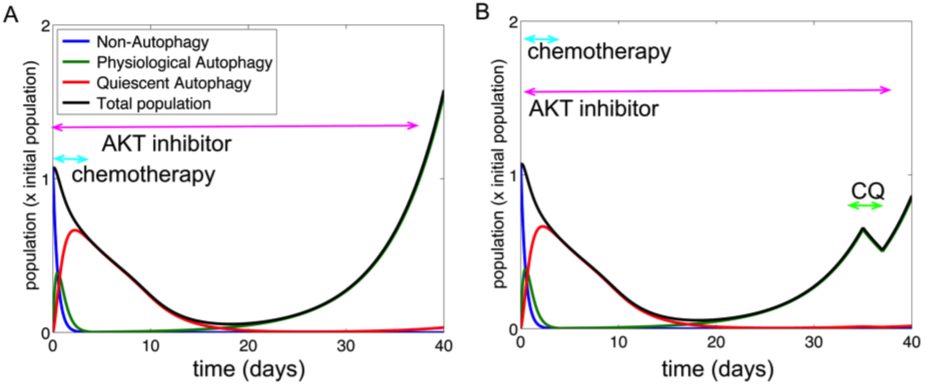
A, 40-day simulation of best combination schedule (#7 in Fig. 2B). B. Addition of CQ on day 30 significantly improves response.

## III. QUICK GUIDE TO THE METHODS (1 PAGE)

### A. Model development

Compartment models are often used to describe transport of material in biological systems. Our compartment model contains three compartments, each containing a well mixed cell population with different autophagy phenotypes. In Fig. 1, boxes represent compartments and arrows represent the connections between the compartments. Every compartment has a number of connections leading to the box and a number of arrows leading from the box. Cells can either flow from one compartment to another, be added to themselves (growth) or they can be removed by death.

In our model (Fig. 1), non-autophagy melanoma cells proliferate (rate: *g*_*N*_), and following treatment can acquire either a physiological autophagy phenotype (rate: *a*_*P*_) or a quiescent autophagy phenotype (rate: *b*_*Q*_). Physiological autophagy cells grow (rate: *g*_*P*_) and can revert to non-autophagy cells (rate: *r*_*P*_), or enter a quiescent/senescent state (rate: *q*_*P*_). Tumor cells having the quiescent autophagy phenotype do not divide, yet these cells can reacquire a physiological autophagy phenotype (rate: *r*_*Q*_) or a non-autophagy phenotype (rate: *r*_*N*_) state. Cells in each compartment die at a fixed rate (*d*_*N,P,Q*_). To model the increased cell death observed in our experimental data, on days 6-9, we included a delay in the model - specifically for the cell death of quiescent autophagy cells (rate: *τ*).

The effects of chemo, AKTi and their combination were also incorporated into the model. As our cell culture experiments showed that chemo triggered cell death with negligible effects on autophagy [1], it was assumed that chemo only augmented cell death. Further, as chemo is effective only in proliferating melanoma populations, we assumed that the therapy increases the death rate of the two proliferating phenotypes, non-autophagy (*d*_*N*_) and physiological autophagy cells (*d*_*P*_). We also assumed that the frequency with which cells became quiescent (*q*_*P*_) increased with chemo. *In vitro* studies showed that while AKTi did not augment cell deaths or effectively inhibit melanoma cell growth, it induced autophagy [1]. Therefore, we assumed that AKTi increases the rate of transitioning to the autophagy phenotypes, *a*_*P*_ and *b*_*Q*_. As combination therapy does not augment cell death compared with chemo, nor significantly increase autophagy relative to AKTi, the combination of the two treatments was modeled by adding the effects of chemo and AKTi. Finally, it was assumed that cell which switch compartment under treatment can only revert back to their original states after treatment is removed. The schematic representation of this compartment converts readily into a system of ordinary differential equations:

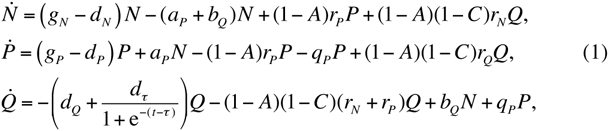

where *d*_*N*_ = *d*_0_ + *c*_*N*_, *d*_*P*_ = *d*_0_ + *c*_*P*_*C*, *q*_*P*_ = *q*_0_ + *c*_*Q*_*C*, *a*_*P*_ = *a*_0_ + *a*_*N*_*A*, and *b*_*Q*_ = *b*_0_ + *b*_*P*_*A*. In equation (1), *A* and *C* are defined by

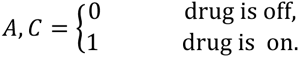

### B. Parameter estimation

Both initial cell populations and the growth rate of non-autophagy cells (*g*_*N*_) are estimated by assuming an exponential growth of untreated cells and finding both the initial value and exponent of the best-fit curve. We used an optimization algorithm called implicit filtering [7] to determine the best remaining parameter set *H* (except *g*_*N*_) that minimized the difference between the predicted number of cells (*N*_*P*_) and experimental results (*N*_*E*_) in 4 conditions: no treatment (*n*); chemo (*c*); AKTi (*a);* and combination (*m*). The mathematical definition of our problem is:

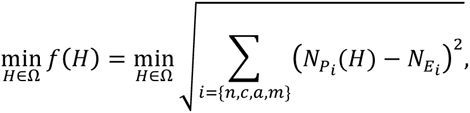

where the goal is to minimize the objective function *f* subject to the condition that *H* ∈ ℝ^*N*^ is in the feasible region:

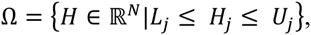

where and *L*_*j*_ and *U*_*j*_ are the upper and lower bound on the *j*th component *H*_*j*_ of the vector *H*.

### C. Other applications

The Compartment modeling approach assumes that material in the compartment is well mixed and homogeneous. The approach can’t capture spatial dynamics, individual diversity (heterogeneity), or stochastic effects. However, the approach is computational efficient, can describe complex systems (interactions) in a simple way, and is easy to formulate. It is is especially useful to model interactions between different states of homogeneous compartments. In cancer modeling field, many compartment models have been developed to model tumor progression [8-17], tumor-stromal interactions [18], tumor-immune interactions[16, 17, 19-32], drug resistance [2, 33], and drug distribution in the body [34]. This approach can also model other biological systems (e.g. ecological) where the material is energy (food) and the compartments represent different species of animals and plants [35]. It has also been widely used to model spread of infectious disease, where compartments represent health status with respect to the pathogen in the system (e.g., susceptible, infectious or recovered) [36, 37].

